# Mining functional genes and characterizing cellular transcriptomic profiles in the single-cell atlas of adult *Spodoptera litura* ovary

**DOI:** 10.64898/2026.03.25.713648

**Authors:** Zhipeng Sun, Liwei Jiang, Xue Dong, Xin Yi, Todd G Nystul, Guohua Zhong

## Abstract

Understanding the reproductive biology of non-model organisms remains challenging due to the limited availability of high-resolution molecular resources. Here, we present a comprehensive single-cell transcriptomic atlas of the adult ovary of *Spodoptera litura* (*S. litura*), a highly polyphagous agricultural pest with a polytrophic meroistuc ovary. By integrating single-cell RNA sequencing with cross-species comparison to *Drosophila melanogaster* (*D. melanogaster*), we define major germline and somatic cell populations and delineate conserved and species-specific features of ovarian cell composition. To enhance the interpretability and reuse of this dataset, we combine transcriptomic profiling with *in situ* hybridization to validate cluster-specific molecular markers across ovarian cell types. We further apply RNA interference (RNAi) to assess the contributions of germline-enriched genes (Hsc70-4, Wech, Polo, Path) to ovarian development and fecundity. Trajectory inference, together with SCENIC and CellChat analyses, provides a system-level view of transcriptional regulatory programs and predicted intercellular communication pathways during oogenesis in *S. litura*. Collectively, this work establishes a valuable resource for studying lepidopteran insect oogenesis, offering a comparative framework for reproductive biology in non-model insects and highlighting potential targets for RNAi-based pest control strategies.

## 1. Introduction

The tobacco cutworm, *Spodoptera litura* (Lepidoptera, Noctuidae), is a highly destructive pest that threatens crops worldwide (1), including cotton, corn, and vegetables (2). Its remarkable reproductive capacity, with females laying thousands of eggs during their short lifespan, contributes to rapid population growth and persistent infestations, exacerbating the challenge of pest control (3, 4). Despite extensive studies on the behavior and ecological effects of this pest, the molecular mechanisms regulating its reproductive development remain poorly understood. As the crucial biological process for their robust reproductive capacity, oogenesis in adult *S. litura* occurs in a polytrophic meroistic ovary, where germline cells interact with somatic cells in a complex and tightly regulated manner through three critical stages: previtellogenesis, vitellogenesis, and choriogenesis (5). Similar to the anatomical arrangement in *Pieris napi*, *Bombyx mori* and *Drosophila melanogaster* (*D. melanogaster*) ovaries, the oocytes of *S. litura* are connected closely to adjacent supporting nurse cells by cytoplasmic junctions. Through a series of precise and intricate physiological progresses, the oocytes ultimately develop into mature eggs and awaits fertilization (6–8).

Extensive studies demonstrate that *S. litura* has evolved marked resistance to most conventional synthetic pesticides such as indoxacarb, neonicotinoid, diamide and even some *Bacillus thuringiensis* toxins (9–12). Apart from this, the widespread application of these synthetic chemicals often nonselectively eliminate beneficial natural enemies, thereby aggravating ecological imbalances as well, which makes it an urgent issue to develop efficient and environmentally-friendly strategies for pest management. In recent years, RNA interference (RNAi)-based pest management has emerged as a promising alternative (13–15). Researchers have employed double-stranded RNA (dsRNA) targeting reproduction-related genes to disrupt growth and development of reproductive system in pests. The spray of nanocarrier-delivered dsRNA of conserved ecdysone receptor and juvenile hormone receptor components exhibited effective inhibition on oviposition and ovarian development in the application of pest management for *Spodoptera frugiperda* and *Adelphocoris suturalis* (16, 17). These cases have demonstrated considerable efficacy of the RNAi technique in reducing population density and field applications. Besides, several recent studies have focused on the integration of transcriptomics and RNAi to identify specific functional genes and microRNAs associated with ovary development and oogenesis in ovary tissues through different pest species such as *Plutella xylostella*, *Nilaparvata lugens* and *Bactrocera dorsalis* (18–20). However, to date the functional characterization of reproduction-related genes in *S. litura* remains unexplored, which largely hinders our understanding of its reproductive biology.

Traditional approaches to study insect ovarian development have largely relied on morphological analysis and histological methods, which provide limited insight into the dynamic cellular processes of tissues formation. The advent of scRNA-seq has revolutionized our understanding of cellular heterogeneity, cell states and cell-cell communication in diverse organism tissues at single-cell resolution (21–23). Nevertheless, no studies have reported the transcriptomic signatures of distinct cell types in *S. litura* ovary. In this study, we present a single-cell transcriptomic atlas of *S. litura* ovary, providing an unprecedented view of its cellular diversity and transcriptional landscapes. By integrating scRNA-seq with *in situ* hybridization and RNAi technique, we identify key molecular markers involved in ovarian development and validate their functional roles. Through a cross-species comparison with *Drosophila melanogaster*, we highlight conserved and unique features of ovarian cell types and developmental trajectories. Our study also explores the transcriptional regulatory networks and intercellular communication pathways that regulate germline-somatic interactions in ovary. This work provides a pivotal resource for uncovering single-cell transcriptomic characteristics of *S. litura* ovary and elucidates unique gene regulatory networks in non-model insects.

## 2. Materials and methods

### 2.1 Insect culture

*S. litura* larvae were reared in climate-controlled incubator under the optimal photoperiod of 14 hours light and 10 hours darkness with a temperature of 26 ± 1℃, and a relative humidity of 50-60%. Larvae were housed in rearing boxes measuring 30 cm×20 cm×10 cm, with fresh feed provided daily. After pupation and emergence, the adult moths were transferred to paper tubes made from folded A4 sheets (80 g weight) and supplied with a 10% honey-water solution.

### 2.2 Preparation of single-cell suspension and sequencing

Ovarian tissues were collected from 48-hour-old adult *S. litura* moths and dissected in Grace’s Insect Basal Medium supplemented with 15% fetal bovine serum (FBS). To optimize the dissociation of the tissue, we used a lysis buffer containing 0.5% Type I collagenase (Invitrogen, cat. no. 17018029) and 1% trypsin (1:250 in EBSS, Invitrogen, cat. no. 27250-018) to incubate for 30-40 minutes with gentle shaking. The dissociation process was carefully monitored to minimize damage to delicate cells, and it was terminated by adding 4 mL of S-FBS medium. The dissociated cell suspension was filtered twice through a sterile 40 μm cell filter. To maximize cell recovery, the centrifuge tube was rinsed 3 times with S-FBS medium. The filtered cell suspension was centrifuged at 500 g, 4°C for 10 minutes. After discarding the supernatant, the cell pelleted were resuspended in 200 μL of S-FBS or EBSS solution. The cell viability was examined by using 0.4% trypan blue (Solarbio, cat. no. T8070) at a proportion of 9:1, and counted by Countess® II Automated Cell Counter. The cell suspension, with a concentration exceeding 1000 cells/μL, and a viability greater than 90%, was processed with Chromium Single-Cell 3’ Library (v2) kit. The procedure included end repair, A-tailing, adaptor ligation, and PCR amplification following the manufacturer’s protocol.

The Cell Ranger Single Cell software package was used to perform quality control on the sequencing data. Before gene alignment, we initially conducted a comparative analysis of the gene similarities between the *S. litura* genome and the *D. melanogaster* genome (Supplementary Data). Then, reads were aligned to the integrated reference genome of *S. litura*genome (https://www.ncbi.nlm.nih.gov/genome/?term=Spodoptera+litura) and *D. melanogaster* genome (https://www.ncbi.nlm.nih.gov/assembly/GCF_000001215.4#/st). After gene alignment, effective reads and unique molecular identifiers (UMIs) were filtered. Data normalization was conducted using Cell Ranger 2.0.0 and Seurat v4.0.4 software.

Quality control for scRNA-seq data was performed based on the following criteria: (1) cells with 200-6000 detected genes were retained; (2) the total number of UMIs per cell was limited to less than 10,000; and (3) the proportion of mitochondrial gene UMIs per cell was restricted to below 30%. Among the initially screened 4915 high-quality cells, the average number of UMIs per cell was 4003, with a filtered mitochondrial UMI proportion of 17.47%.

### 2.3 Clustering, cell annotation and marker screening

The clustering analysis of 4915 cells was performed using the single-cell gene expression matrix generated by Cell Ranger. The analysis was carried out with the FindClusters function in the Seurat package, applying a resolution of 0.5, a log fold change threshold of 0.25, and a minimum proportion of 0.01, resulting in a tSNE plot of 14 distinct cell clusters. The characteristic genes for each cluster were identified using the FindAllMarkers function in Seurat. Cell types were then assigned based on the SingleR package and known marker genes. To annotate the 14 clusters, we referenced marker genes according to previous reports in *B. mori* and *D. melanogaster*. The cell types were defined by comparing the expression patterns of these marker genes, enabling a cross-species comparison and characterization of the cell types.

### 2.4 Trajectory analysis

The single-cell developmental trajectory of somatic cell clusters and germline clusters was analyzed using a gene expression matrix in the monocle package (v2.22.0) within R software. This approach generated visualized trajectories, displaying tips and branches in a reduced-dimensional space. Cell types were first annotated using Seurat, and developmental trajectories were constructed using unsupervised analysis methods. The resulting cell trajectory plots were created based on pseudotime values, cell types, and selected genes.

### 2.5 RNA-fluorescence *in situ* hybridization

The *in situ* probes were synthesized by using genome DNA of *S. litura* as a template for PCR amplification with the KAPA HiFi PCR High-Fidelity Enzyme Kit (Roche Diagnostics, cat. no. 07958927001). The upstream primer is preceded by the T7 sequence: TAATACGACTCACTATAGG GAGA, and the downstream primer is preceded by the SP6 sequence: ATTTAGGTGACACTATAGAAGNG. The *in situ* primer sequences were listed in **Table S1**. All procedures were conducted on ice to maintain sample integrity. Amplification products were analyzed via gel electrophoresis, and the target DNA bands were extracted and purified. Sense and antisense digoxigenin probes were synthesized by using DIG RNA Labeling Kit (Roche, cat. no.11175025910) according to instruction. Briefly, the mixture was incubated in a metal bath at 37°C for 2 hours. The DNase I was added, and continuously incubated the mixture for 1 hour. To terminate the reaction, 30 µL of LiCl was added to the tube and thoroughly mixed. Samples were frozen at -20°C for 30 minutes. Then, the samples were centrifuged at 12,000 rpm for 15 minutes, and the supernatant was carefully removed. After adding 1 mL anhydrous ethanol, the sample was centrifuged at 4°C for 10 minutes.

The protocol of *in situ* hybridization was described in our previous work (24), with minor modification to incubation which was conducted at 65°C.

### 2.6 Gene knockdown by dsRNA injection

The dsRNAs targeting specific genes were designed and synthesized using the T7 RiboMAX Express RNAi System (Promega, USA). The sequences for primer pairs were listed in **Table S2**. The green fluorescent protein (GFP) was used as a control. The synthesized dsRNAs were purified using the RNeasy MinElute Cleanup Kit (Qiagen, Germany). To assess the effects of marker genes on development of ovary, 5 μL (5 μg/μL) of dsRNA was injected into the third abdominal segment of newly emerged female moths. A booster injection of the same dsRNA dose was administered in the next day.

### 2.7 Assessment of reproductive capacity in *S. litura*

The average oviposition of *S. litura* was calculated after the second injection for 72 hours. Additionally, adult female moths were dissected to measure the ovarian weight and the lengths of ovarioles at previtellogenesis stage, vitellogenesis stage and choriogenesis stage. The representative morphological images were captured under the stereomicroscope (Leica, S APO).

### 2.8 RT-qPCR detection

Total RNA was extracted from adult tissues with Total RNA Isolation Kit I (OMEGA, cat. no. R6834-00). First-strand cDNA synthesis was performed using Evo M-MLV Reverse Transcriptase Mix Kit (AG, cat. no. AG11728). Real-Time quantitative PCR (RT-qPCR) was carried out with SYBR Premix EX Taq^TM^ Kit (TaKaRa, cat. no. DRR041A) and conducted on CFX96 System (BioRad) following: initial denaturation at 95°C for 30 s, 40 cycles of 95°C for 10 s, annealing at 60°C for 10 s, elongation at 72°C for 30 s. All specific sequences for primer pairs were listed in **Table S3**.

### 2.9 GO/KEGG enrichment analysis

To perform functional annotations, the Gene Ontology (GO) database (http://www.geneontology.org/) was utilized for categorizing differentially expressed genes between somatic cell and germ cell clusters according to cellular component (CC), biological process (BP), and molecular function (MF). Pathway enrichment analysis was conducted using the Kyoto Encyclopedia of Genes and Genomes (KEGG) database (http://www.genome.jp/kegg/). Enrichment of GO terms and KEGG pathways was assessed using Fisher’s exact test. To account for multiple comparisons, the Benjamini-Hochberg correction was applied to adjust the p-values. Only those functional categories and pathways with adjusted p-values below 0.05 were considered statistically significant.

### 2.10 The regulatory network analysis

To build a transcription factor (TF) regulatory network from our scRNA-seq data, we employed the SCENIC package (version 1.2.4). SCENIC analysis was performed on all individual cells to assess variations in TF activity and their downstream target genes across different somatic cell clusters and germline clusters. The TF activity quantification matrix was incorporated into the Seurat object, and the activity of each regulon was evaluated using Area Under the Curve (AUC) scores with the AUCell R package.

### 2.11 Cell-cell communication analysis

Cell-cell signaling interactions were analyzed using CellChat V2.1.0 software to construct a ligand-receptor interaction network. The single-cell gene expression matrix and cell grouping data were provided as input, while the ligand pairing information from the CellChatDB database was used to evaluate communication strength between cell clusters. Using default settings, we calculated the communication probability at the signaling pathway level by evaluating ligand-receptor interactions within each pathway. The significance of the ligand-receptor pairs in key signaling pathways was visualized using a dot plot.

### 2.12 Cross-species cell type comparison

The Sankey plot was generated using the networkD3 package and visualized through the sankeyNetwork function. To assess the similarity between different cell types in *D. melanogaster* and *S. litura*, Pearson’s correlation coefficient was calculated using the R package. Statistical significance was determined with a p-value threshold of ≤0.05.

### 2.13 Statistical analysis

The statistical analyses were conducted by Wilcoxon rank sum test or one-way analysis of variance (ANOVA) followed by Duncan’s Multiple Range Test (DMRT), which performed via the SAS statistical software package version 8.1 (Microsoft, USA). Graphical images were generated using GraphPad Prism software package version 9.5 for windows (Microsoft, USA).

## 3. Results

### 4.1. Cell type prediction in *S. litura* ovary through cross-species analysis

Initially, we intended to collect the parts of ovary at previtellogenesis and vitellogenesis stages to perform single-cell RNA sequence, since the cell strainer only allows cell with a maximum diameter of 40 μm to pass through. A barcoded cell suspension was converted into a sequencing library, and processed using the 10× Chromium single-cell RNA sequencing platform (Figure 1A). After quality control, 4,915 high-quality cells were retained from an initial pool of 12,623 viable cells.

**Fig. 1.**
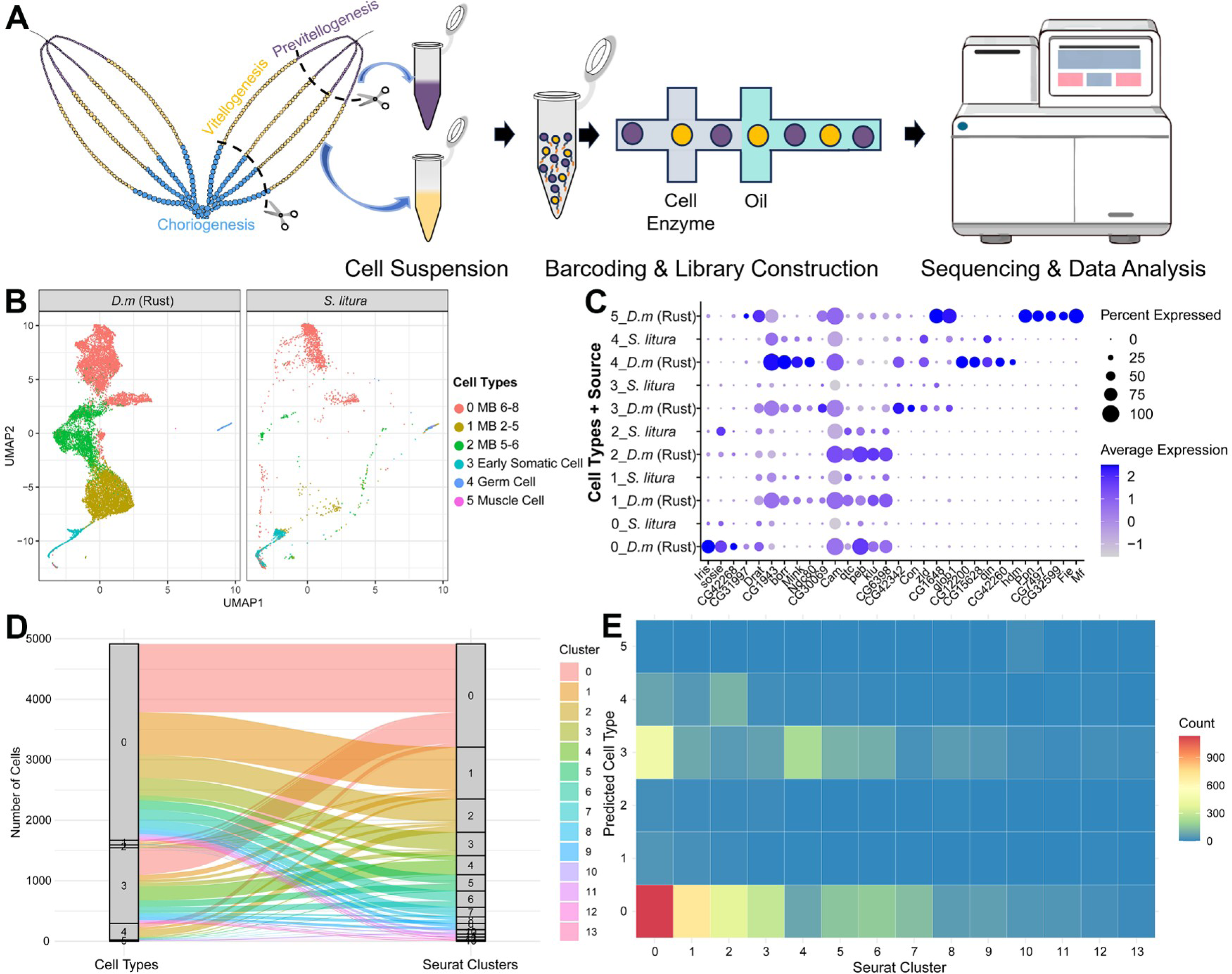
Cross-species analysis of ovarian cells between *D. melanogaster* and *S. litura*. **A.** Schematic workflow of sample preparation and scRNA sequencing. Ovarian tissues at the previtellogenesis and the vitellogenesis stages were collected for scRNA-seq. **B.** UMAP plots of referenced *Drosophila* ovary dataset and *S. litura* ovary dataset. **C.** Gene expression profiles of selected markers in distinct cell types. **D.** The sankey plot visualizes the conserved relationships and divergences between 5 predicted cell types and 14 cell clusters in *S. litura*. **E.** Heatmap for the abundance of count number between 5 predicted cell types and 14 cell clusters in *S. litura*.

Although *S. litura* and *D. melanogaster* belong to different insect orders, both species possess polytrophic meroistic ovaries which suggests the possibility of conserved cellular compositions and developmental mechanisms between the two species. Therefore, we speculated that the ovarian cell types in *S. litura* may exhibit conservation with those in *D. melanogaster*. An efficient application for cell type annotation is using known cell types in a model specie to label the cell types in a non-model organism. We firstly referenced the recent published data and identified the atlas of *Drosophila* ovary by using specific marker (Figure S1A and S1B). Then, visualized our *S. litura* dataset and *D. melanogaster* dataset with dimensionality reduction using uniform manifold approximation and projection (UMAP) and grouped cells into 6 distinct cell types with high prediction confidence (Figure 1B and Figure S1C). The gene expression profiles of marker genes in 6 predicted cell types exhibited a similarity between *S. litura* dataset and *D. melanogaster* dataset (Figure 1C). Further, the result of sankey diagram revealed that 14 seurat clusters of *S. litura* dataset mapped to 5 cell types (Figure 1D). By calculating the enrichment of cell counts among 14 seurat clusters, we concluded that *S. litura* ovary mainly consist of main body follicle cell, germ cell and early somatic cell (Figure 1E).

### 4.2. Single-cell RNA landscape of cell-type classification in adult *S. litura* ovary

The adult ovary of *S. litura* comprises three primary regions: previtellogenesis, vitellogenesis, and choriogenesis. In this study, the vitellogenic region was further subdivided into anterior, central, and posterior segments based on distinct histomorphological features such as chamber color, size, and shape (Figure 2A). Subsequently, dimensionality reduction and clustering were performed using the t-distributed stochastic neighbor embedding (tSNE) method in conjunction with the Seurat algorithm at a default resolution (R = 0.5), resulting in 14 distinct clusters (Figure 2B).

**Fig. 2.**
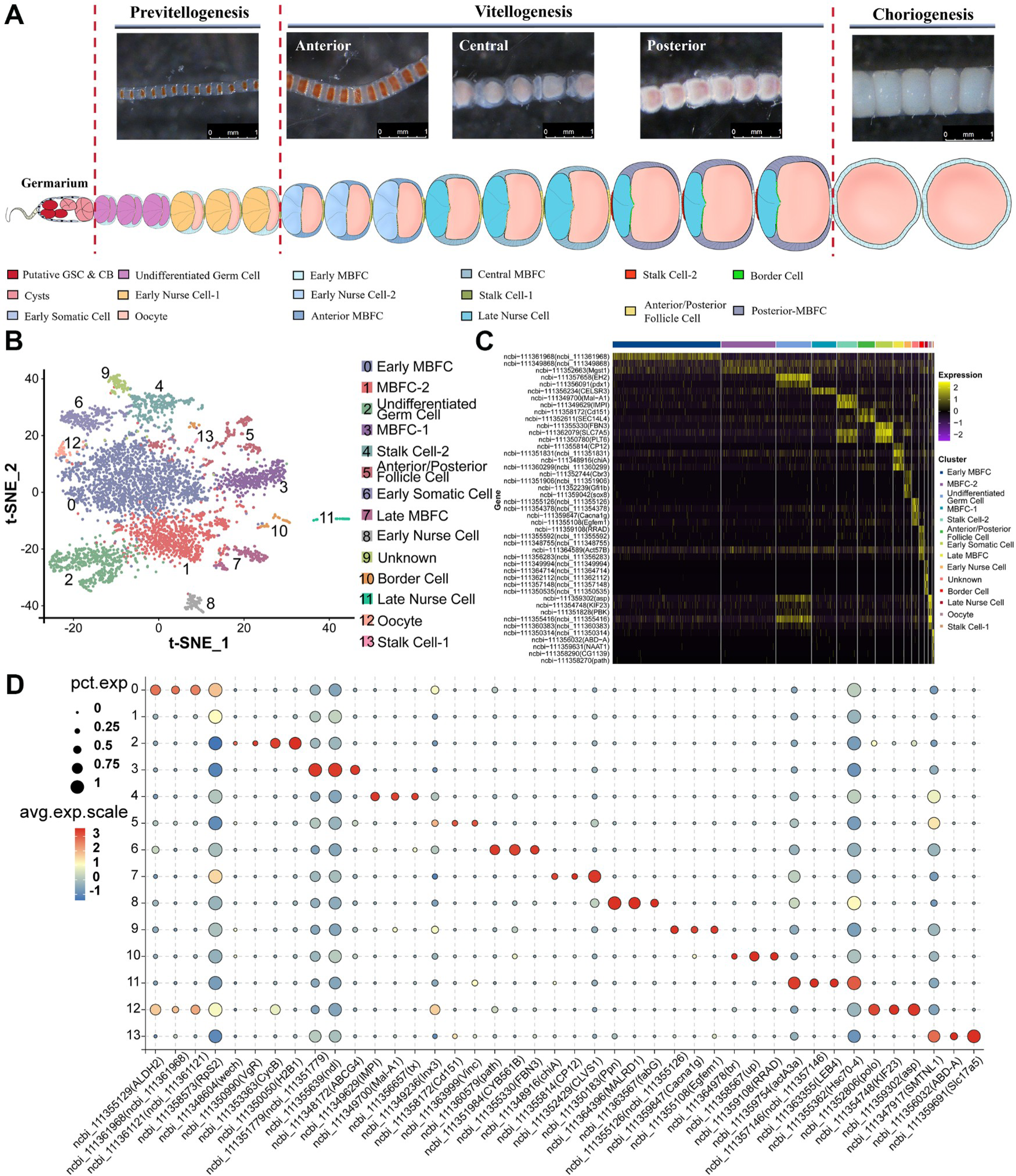
Single-cell transcriptomic classification of cell types in *S. litura* ovary. **A.** An illustration of single ovariole coupling with representative images during previtellogenesis, vitellogenesis and choriogenesis in *S. litura* adult ovary. Partial ovariole at vitellogenesis is morphologically divided into anterior, central and posterior vitellogenesis. **B.** A t-SNE projection of 14 identified cell clusters in our dataset. **C.** Heatmap of top marker genes in clusters. The gene ID and its homologous gene name were listed on the left, and each column represents an individual cell. **D.** Dot plot of selected specific genes showing the scaled expression and percentage in each inferred cell cluster. The intensity of the color is positively correlated with gene expression levels, and the red color shows a higher average gene expression, while the light color shows a lower average gene expression.

Given the absence of well-established marker genes for *S. litura* ovarian cells, we assigned cell type identities by referencing conserved orthologous markers characterized in *D. melanogaster*. For example, follicle cells were identified using *ndl* (ncbi_111355639) and *Br* (ncbi_111364978), germ cells were defined by *CycB* (ncbi_111353363), *polo* (ncbi_111352806), *KIF23* (ncbi_111354748), and *Wech* (ncbi_111348604), and oocytes were recognized using *VgR* (ncbi_111350990) (25,26) (Figure 2C). Some of these annotations were corroborated through follow-up experiments. Meanwhile, we also selected 3-4 cluster-specific marker genes from top 10 enriched genes in each cluster as the candidates for *in situ* hybridization validation (Figure 1E). Finally, Early Somatic Cells (5.47%), Early MBFC (34.73%), MBFC-1 (7.85%), MBFC-2 (17.44%), Anterior/Posterior Follicle Cells (5.47%), Late MBFC (3.23%), Border Cells (1.49%), Stalk Cell-1 (0.43%), Stalk Cell-2 (6.41%), Undifferentiated Germ Cells (11.21%), Early Nurse Cells (2.22%), Late Nurse Cells (1%), Oocytes (0.98%), and an Unknown cluster (2.08%) were identified in *S. litura* ovary (Figure 2D).

### 4.2. Dynamic gene expression profiles of germline-specific marker

There were 6 cluster-specific high expression genes facilitating to distinguish 4 germline clusters and 1 early somatic cell cluster (Figure 3A). The *Cyclin B* (*cycB*) is particularly involved in the control of the transition from the G2 phase to the M phase (mitosis) of the cell cycle in the processes of embryogenesis and cellular differentiation in insects. The outcomes of RNA *in situ* hybridization shows that *CycB* was expressed in the early germ cells at early previtellogenesis stage, which is recognized as the undifferentiated germ cell cluster. Meanwhile, *path*, a specific marker for early somatic cells, was widely detected during early previtellogenesis (Figure 3B). The expression of *PPn* was detected in early nurse cells during the previtellogenesis, and presented in anterior region of the vitellogenic ovary (Figure 3C). At the central and posterior vitellogenic region, *actA3a* was highly expressed in late nurse cells (Figure 3D). Interestingly, although the expression profile of *wech* was restrict to the undifferentiated germ cell cluster in our scRNA data, we found that the expression of *wech* was detectable in oocyte through previtellogenesis and vitellogenesis (Figure 3E). *Polo*, a marker of oocyte cluster, was validated in the oocyte at previtellogenesis and anterior/central regions of vitellogenesis (Figure 3E). Cell trajectory analysis of 4 germline clusters revealed that undifferentiated germ cell, serving as the origin point of differentiation, will gradually diverge into early/late nurse cell and oocyte during oogenesis. Consistent with the results of *in situ* experiments, pseudotime analysis assigning with marker genes showed dynamic expression patterns of multiple markers along the germ cell differentiation trajectory (Figure 3F).

**Fig. 3.**
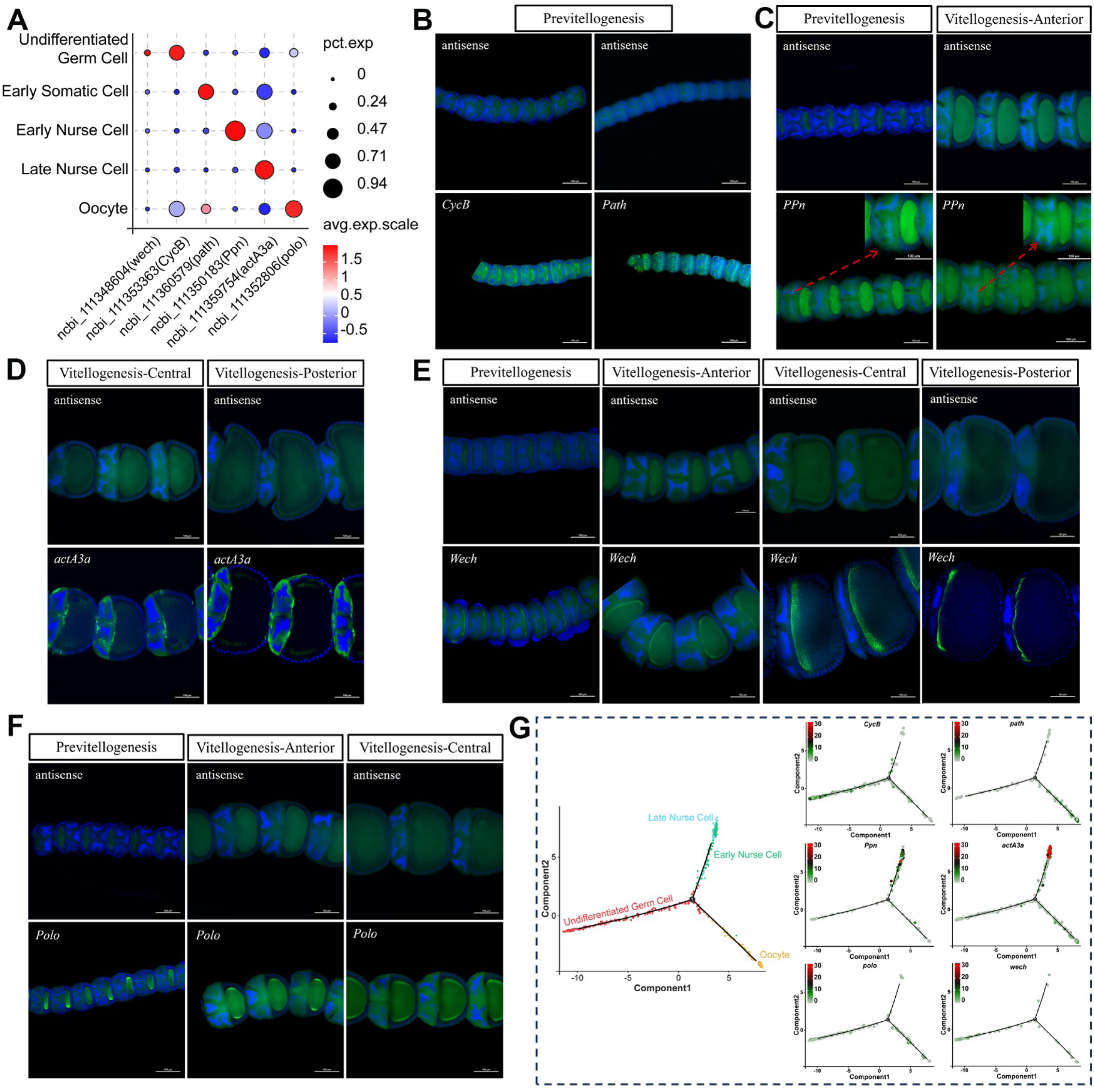
Identification of 4 germ cell clusters and *path*-positive early somatic cell cluster detecting by *in situ* hybridization. **A.** Dot plot shows the profile of 5 candidate marker genes among germline clusters. The dot color represents average expression level of marker gene and dot size represents percentage of cells within each inferred cell cluster expressing that gene. **B-F.** Confocal stack images of marker gene expression patterns (ncbi_111348604 (*wech*), ncbi_111353363 (*CycB*), ncbi_111360579 (*path*), ncbi_111350183 (*Ppn*), ncbi_111359754 (*actA3a*) and ncbi_111352806 (*polo*)) detecting by whole-mount RNA *in situ* hybridization together with the negative control respectively during previtellogenesis and vitellogenesis. Several images were magnified in part. Red arrow points out the region of *in situ* signals. The green fluorescence signals highlight the expression of target genes. Scale bar is 100 μm. **G.** Differentiation trajectory construction with Undifferentiated Germ Cell (red), Early Nurse Cell (green), Oocyte (orange) and Late Nurse Cell (blue) and pseudotime trajectory of marker genes in these clusters.

### 4.3. Molecular markers for the characterization of somatic clusters

Assigning cell types to somatic cell clusters is more challenging than identifying germline clusters, as some genes were expressed across multiple clusters. To assign identities for different somatic clusters, we screened out several marker genes which are highly expressed or upregulated among somatic clusters in our scRNA data (Figure 4A) and assessed their mRNA expression patterns in *S. litura* ovary. Notably, *ALDH2*, a mitochondrial isoform of aldehyde dehydrogenase, was enriched in the oocyte at previtillogenesis, but also presented in early MBFC at anterior region of vitellogenesis, suggesting that *ALDH2* may be involved in the cell-cell interaction between oocyte and early MBFC (Figure 4B). According to the order of spatial and temporal expression pattern, we identified that MBFC-1 cells expressed titin throughout vitellogenesis stage (Figure 4D), and MBFC-2 cells specifically expressed *RpS2* gene in the central and posterior regions of vitellogenesis (Figure 4C), while late MBFC cells weakly expressed *CLVS1* only in the posterior region of vitellogenesis (Figure 4E). *Inx3* was used to visualize the anterior/posterior follicle cells population. We observed that follicle cells expressing *Inx3* were located on the boundary between nurse cell and oocyte at previtillogenesis and anterior/central regions of vitellogenesis (Figure 4F).

**Fig. 4.**
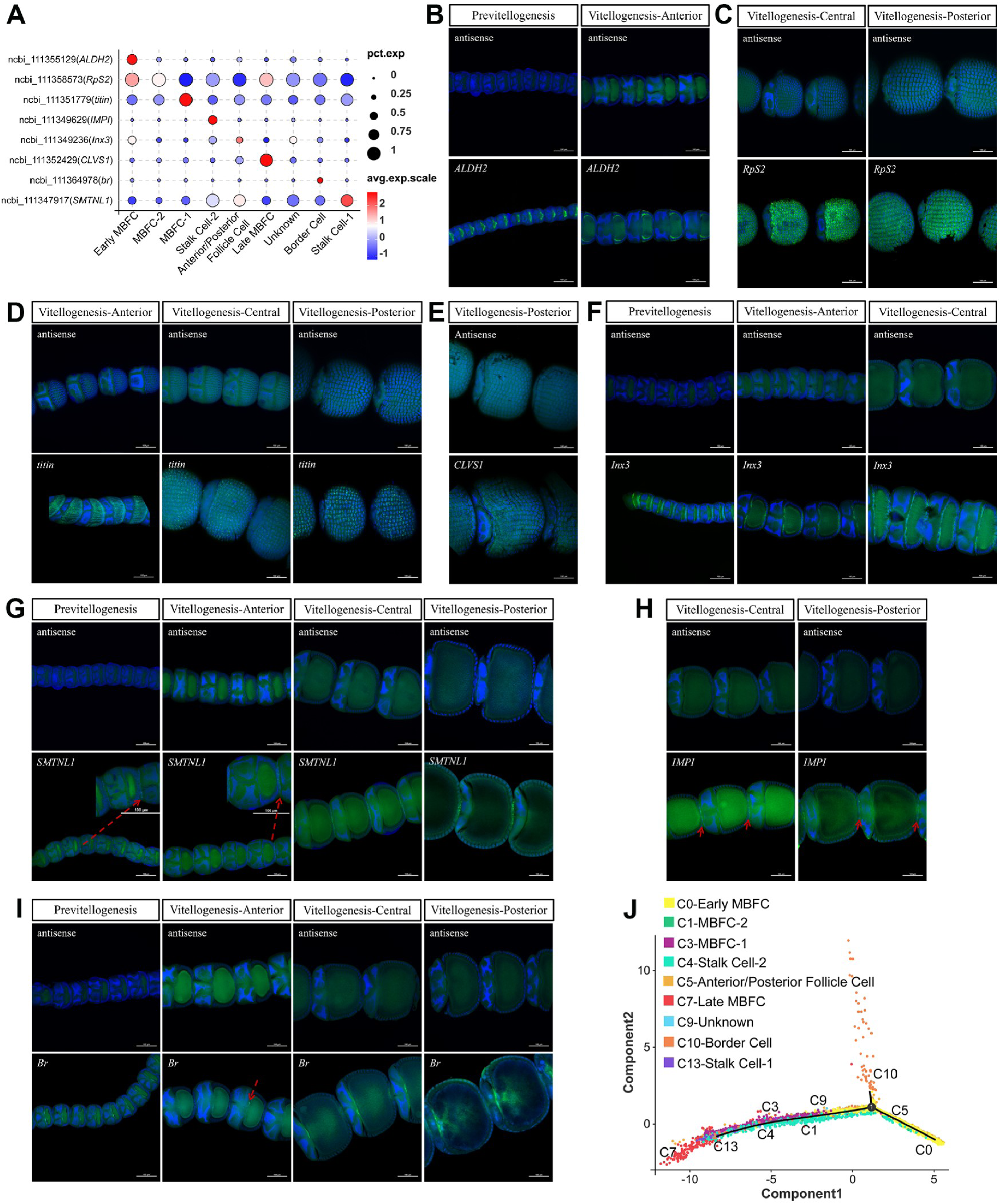
Expression patterns of marker genes in different cell types of somatic cells. **A.** Dot plot showing the selected marker genes distinguishing different somatic cell clusters. **B-I.** Confocal stack images of marker gene expression patterns (ncbi_111355129 (*ALDH2*), ncbi_111358573 (*RpS2*), ncbi_111351779 (*titin*), ncbi_111349629 (*IMPI*), ncbi_111349236 (*Inx3*), ncbi_111352429 (*CLVS1*), ncbi_111364978 (*br*) and ncbi_111347917 (*SMTNL1*)) were obtained through whole-mount RNA *in situ* hybridization, along with the negative control respectively during previtellogenesis and vitellogenesis. Several images were magnified in part. Red arrow points out the region of *in situ* signals. The green fluorescence signals highlight the expression of target genes. Scale bar is 100 μm. **J.** Pseudotime trajectory showing the potential differentiation orientation of different somatic cell types from early MBFC to late MBFC.

To assign the identity of stalk cell subtype, we labeled *SMTNL1* and *IMPI* in *S. litura* ovary, and found that stalk cell-1 strongly expressed *SMTNL1* at previtillogenesis and vitellogenesis stages (Figure 4G), while stalk cell-2 specifically expressed *IMPI* at central/posterior regions of vitellogenesis (Figure 4H). The border cells interact with surrounding follicle cells and the oocyte through cell-cell communication, influencing the polarization and position of oocyte during egg chamber development. We used *Br* to visualize the anatomical position of border cell in egg chamber during previtillogenesis and vitellogenesis (Figure 4I). However, one unknown cluster was undetectable due to the lack of effective makers. To further characterize the cellular types and potential differential relationship of somatic clusters, we performed monocle analysis on newly validated cell clusters, and found that all kinds of follicle cell and stalk cells were fitted well onto the same branch in the pseudotime trajectory except for the border cell suggesting that the follicle cells and stalk cells may exhibit certain interactions during the development of the egg chamber (Figure 4J).

### 4.4. RNAi-based functional analysis of marker genes involving in the development of *S. litura* ovary

To explore the potential function of germline-related genes, dsRNA interference technology is a favorable approach to silence the target genes in insects. We first quantify the gene expression level in ovary, head, fat body and abdomen of adult *S. litura* by RT-qPCR. The result showed that the expression of *Hsc70-4*, *Wech*, *PPn*, *Polo* and *Path* were significantly enriched in ovaries tissue, demonstrating that these genes ought to mainly function in *S. litura* ovary (Figure 5A). After dsRNA injection, gene expression levels of target genes were significantly downregulated, suggesting a high RNAi efficiency (Figure 5B). Following two consecutive days of dsRNA injection, we found that the ovarian weight in RNAi-treated adult moth was significantly decreased comparing with that of in control group (Figure 5C). In addition, we observed the dramatically morphological changes on RNAi-treated ovaries (Figure 5D) and examined the regional length of treated ovary at previtillogenesis, vitellogenesis and choriogenesis. The result indicated that the ovarian length at choriogenesis was significantly decreased in RNAi-treated group, while the ovarian length at vitellogenesis was significantly increased after *wech*- and *PPn*-RNAi, suggesting that the silence of *wech* and *PPn* may arrest the development of germ cells in vitellogenesis to certain extent (Figure 5E). Subsequently, we also found that the fecundity was strikingly reduced by calculating the cumulative number of eggs within 72 hours after RNAi (Figure 5F). These results reveal that germline-specific genes play a crucial role in the development of egg mature.

**Fig. 5.**
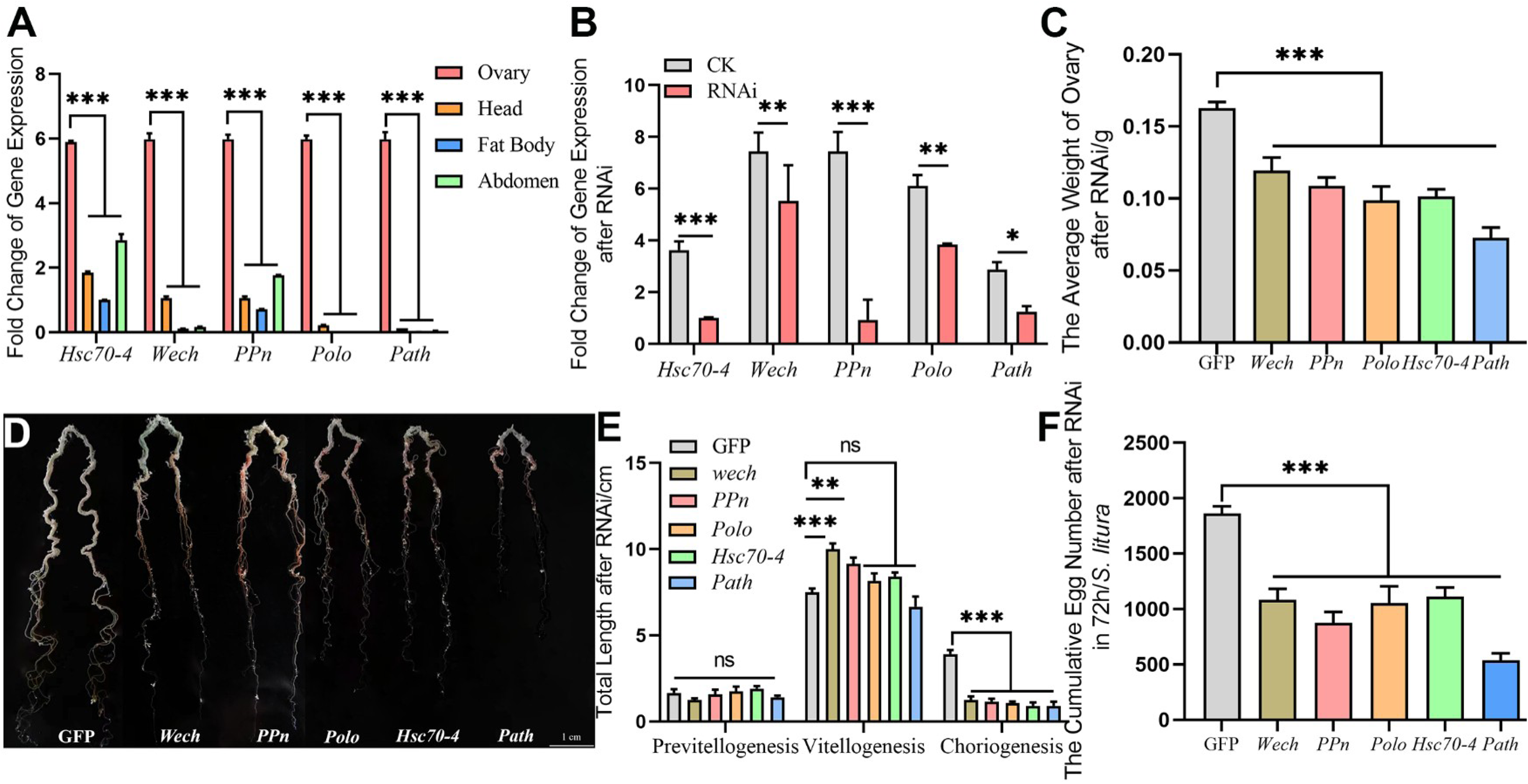
Functional validation of germline-specific genes. **A.** Relative expression level of germ cell marker genes (*Hsc70-4*, *Wech*, *PPn*, *Polo*, *Path*) among ovary, head, fat body and abdomen. **B.** Analysis of relative expression level of selected genes in ovaries after dsRNA injection. **C.** Analysis of average weight of *S. litura* adult ovary after RNAi. **D.** Observation of morphological changes on whole *S. litura* ovary after RNAi. **E.** Analysis of total length of ovaries during previtellogenesis, vitellogenesis and choriogenesis. **F.** Quantification of cumulative egg number after RNAi within 72 h. Wilcoxon rank sum test is used as statistical analysis. All data are showed as mean ± standard deviation, and ns indicates no significant difference (*p* > 0.05), \**p* ≤ 0.05, \*\**p* ≤ 0.01 and \*\*\**p* ≤ 0.001.

### 4.5. Transcriptomic feature analysis of germ cells and somatic cells

To thoroughly depict the signature of transcriptional expression in germ cells and somatic cells, we performed GO analysis and KEGG pathway enrichment analysis on the differentially expressed genes in somatic cell clusters and germ cell clusters. Go term analysis revealed that the number of differential genes categorized in the term of biological process and molecular function in germ cells was larger than that of in somatic cells, while the number of functional genes involved in cellular component such as cell, cell part and macromolecular complex in somatic cells was larger than that of in germ cell clusters (Figure 6A and Figure S2A-B). Additionally, the germ cell clusters exhibit a higher number of genes in most KEGG pathway categories, particularly in infectious diseases, metabolism, organismal systems, and environmental information processing, suggesting that germ cells may possess more complex and diverse gene regulatory mechanisms in *S. litura* (Figure 6B).

**Fig. 6.**
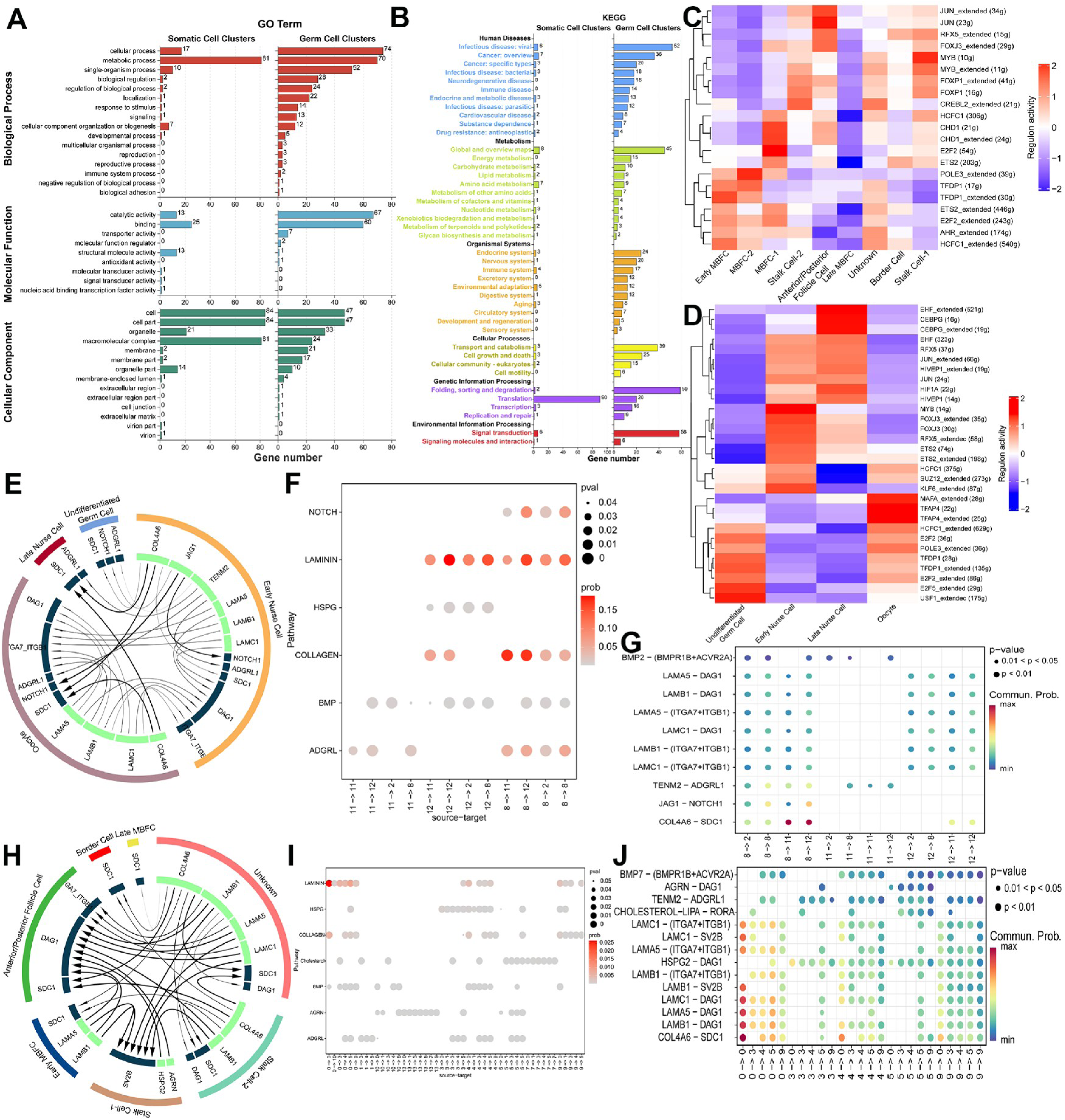
Comparative analysis of transcriptomic features in somatic cells and germline cells in *S. litura*. **A.** Gene Ontology analysis was performed for DEGs in somatic cell clusters and germ cell clusters. The number of DEGs associated with each GO term was noted. **B.** KEGG pathway enrichment analysis showing the number of genes involved in various biological pathways in somatic cell clusters and germ cell clusters. **C-D.** Heatmap showing differences in TF activity across somatic cells (C) and germ cells by using SCENIC analysis. (D). **E.** The network diagram presents the strength of ligand-receptor interactions within and between germ cell clusters. The line with arrowhead indicates the inferred incoming communication pattern in each cluster and the inferred outgoing communication pattern among clusters. **F.** Dot plot shows the top 6 interaction signaling pathway with the lowest p-values and the highest communication probabilities of ligand-receptor pairs among 4 germline clusters. G. Dot plot shows the expression abundance of ligand-receptor pairs with significant communication probabilities (p-value≤0.05), ranked by gene expression levels of ligand and receptor in germline clusters. **H.** The network diagram presents the strength of ligand-receptor interactions within and between somatic cell clusters. **I.** Dot plot shows the top 7 interaction signaling pathway with the lowest p-values and the highest communication probabilities of ligand-receptor pairs among somatic clusters. J. Dot plot shows the expression abundance of ligand-receptor pairs with significant communication probabilities (p-value≤0.05), ranked by gene expression levels of ligand and receptor in germline clusters.

Next, we investigated whether the identified cell types are defined by distinct gene regulatory networks and the activity of specific transcription factor (TF) combinations. SCENIC analysis revealed that early MBFC shared similar active regulons with MBFC-2, while CHD1, CHD_extended and E2F2 were significantly enriched in MBFC-1. Anterior/posterior follicle cell cluster exhibited high activity in JUN_extended, JUN, RFX5_extended and FOXJ3_extended. Notably, most of regulon activity in late MBFC was significantly downregulated (Figure 6C). For germline clusters, E2F5_extended and USF1_extended were specifically enriched in Undifferented Germ Cell cluster, and MAFA_extended, TFAP4 and TFAP4_extended were significantly enriched in oocyte. In two nurse cell clusters, MYB and KLF6_extended were highly enriched in early nurse cell, while EHF_extended, CEBPG and CEBPG_extended were specifically enriched in late nurse cell (Figure 6D). These results indicates that various TF-based regulatory networks are pivotal for cell type specification in *S. litura* ovary.

To deepen our global understanding of cell-cell communication landscape in the *S. litura* ovary, we performed CellChat analysis using CellphoneDB to predict cellular receptor-ligand interactions in somatic cells and germ cells. The result showed that early nurse cell closely interacted with undifferented germ cell, oocyte and late nurse cell with a high interaction strength, and the oocyte was able to give the feedback to early nurse cell (Figure 6E). Two signaling pathways (laminin and collagen) were inferred to actively execute cellular communications for both early nurse cell and oocyte (Figure 6F). A single ligand-receptor pair COL4A6-SDC1 was identified as the most significant singling from early nurse cell to oocyte and late nurse cell, suggesting the importance of this ligand-receptor pair in germ cell interactions (Figure 6G). Among somatic cells, we found that early MBFC and unknown cluster frequently communicated to anterior/posterior follicle cell and stalk cell-1 (Figure 6H). Interestingly, we found that laminin and collagen pathway were also significantly active in early MBFC, and laminin pathway exhibited high activity from early MBFC to MBFC-1, stalk cell-2 and anterior/posterior follicle cell respectively (Figure 5I). Multiple igand-receptor pairs such COL4A6-SDC1, LAMB1-DAG1, LAMBA5-DAG1, LAMC1-DAG1, LAMA5-(ITGA7+ITGB1) and LAMC1-(ITGA7+ITGB1) were identified as high signaling from early MBFC to stalk cell-2 and anterior/posterior follicle cell respectively (Figure 5J).

## 4. Discussion

Insect ovaries provide the cellular and molecular foundation for germline maintenance and reproductive output, yet our understanding of ovarian cell-type organization and regulatory programs remains heavily biased toward a small number of model species. Although single-cell transcriptomics has transfromed the analysis of cellular heterogeneity in diverse tissues, its application to non-model insects has been constrained by limited reference atlases and poorly defined cell-type markers (27–29). In this study, we establish a single-cell transcriptomic resource for *S. litura* ovary, enabling systematic dissection of ovarian cell states and their transcriptional features at single-cell resolution.

*Drosophila* and Many Lepidoptera insects such as *Pieris napi*, *Bombyx mori* and *S. litura* have polytrophic meroistic ovary, suggesting a tissue-specific revolutionary conservation between these two species (6–8). Leveraging the conserved polytrophic meroistic architecture shared between *D. melanogaster* and *S. litura*, we implemented a cross-species orthology-based annotation strategy to map 14 transcriptionally distinct clusters onto 5 predicted cell types and to integrate these findings into a schematic model of the adult *S. litura* ovariole (Figure 2A). Compared with traditional histological and antibody-based approaches (30–32), our single-cell framework resolves ovarian cellular complexity with substantially higher resolution, identifying 9 somatic cell clusters and 4 germline cell clusters in *S. litura* ovary and provideing molecular markers for each cell type. Spatial validation by *in situ* hybridization further substantiates the robustness and interpretability of the atlas. Notably, undifferentiated germ cells were defined by *cycB* expression, however, putative germline stem cells (GSCs) and early germ cysts were not recovered. This likely reflects a low GSC abundance in adult ovaries and the inherent sampling limitions of droplet-based single-cell sequencing. These observations highlight a general challenge for ovarian single-cell studies and underscore the need for future enrichment approaches to capture rare germ cell populations.

It is reported that *S. litura* has developed to strong adaptability and resistance to commercial pesticides, and genetically evolved multi-copy detoxification genes in its genome (1, 33). In contrast to the application of conventional pesticides, RNAi-based pest management is a promising strategy to control agricultural pests with high efficiency and low toxicity to non-targeted organisms (34–36). However, the identity and functional relevance of reproduction-associated genes in *S. litura* remain poorly characterized, limiting the rational selection of effective RNAi targets. Functional interrogation of germline-specific marker genes using RNAi technique demonstrates that knock down of *Hsc70-4*, *Wech*, *PPn*, *Polo* and *Path* markedly impairs ovarian development and reduces fecundity (Figures 5C-5F), indicating that our single-cell atlas constitutes a valuable reference for systematically prioritizing novel reproductive targets in *S. litura* and other lepidopteran pests.

Beyond cell-type annotation, this resouce enables comparative and functional analyses of ovarian regulatory programs. Pathway enrichment analyses reveal that germ cells are characterized by transcriptional programs associated with cancer disease, carbohydrate metabolism, cellular transport and catabolism and DNA replication and repair were enriched in germ cells, mirroring features in previously described *Drosophila* germ cells (25, 37). Moreover, one regulon E2F2_extended that has been identified as an active gene regulatory network in undifferentiated germ cell, older germ cell and escort cells in *Drosophila* ovary also exhibited specific activity in undifferentiated germ cell, oocyte, early MBFC and early somatic cells in *S. litura* ovary, supporting the evolutionary conservation of key regulatory principles governing insect oogenesis. Taken together, our study provides a high-resolution, experimentally validated single-cell resource for the *S. litura* ovary and establishes a comparative framework for investigating conserved and species-specific mechanisms of insect oogenesis, with broad utility for reproductive biology and functional genomics in non-model insects.

## 5. Limitations of the study

Our present work identified a useful atlas of single-cell *S. litura* ovary transcriptomics combing with SCENIC and CellChat analysis, which provides essential resources for grasping the heterogeneity and cellular regulatory networks in *S. litura* ovary. Despite the comprehensive dataset generated in this study, one limitation is the difficulty in capturing GSCs, which are scarce in ovary tissues. Meanwhile, some limitations including sample size and sequencing depth may hinder a comprehensive understanding of complex intercellular interactions in *S. litura* ovary. Future studies should focus on developing improved techniques for GSCs enrichment or combining with more refined cell isolation methods such as fluorescence-activated cell sorting (FACS) or microfluidic-based approaches. Moreover, integrating multi-omics technologies such as proteomics, epigenomics, and spatial transcriptomics could offer a more holistic understanding of gene regulation and cellular interactions in *S. litura* ovaries.

## Data availability

The raw sequence data reported in this paper have been deposited in the Genome Sequence Archive (Genomics, Proteomics & Bioinformatics 2021) in National Genomics Data Center (Nucleic Acids Res 2022), China National Center for Bioinformation / Beijing Institute of Genomics, Chinese Academy of Sciences (GSA: CRA030536) that are publicly accessible at https://ngdc.cncb.ac.cn/gsa. The referenced datasets in this study are available in GEO databases (GSE210822)

## Acknowledgments

This study was supported by National Natural Science Foundation of China (Grant No. 32500403). We gratefully thank Guangzhou Genedenovo Biotechnology Co., Ltd for the assistance in sequencing and bioinformatics analysis.

## Author Contributions

Zhipeng Sun and Liwei Jiang designed and performed all experiments and validation. Xue Dong contributed to the sample collection, visualization and data analysis. Zhipeng Sun wrote original draft and revised the manuscript. Xin Yi provided professional guidance and review the manuscript. Guohua Zhong and Todd G Nystul supervised this research and review the manuscript.

## Supplementary Figures and Tables

Table S1. The primer sequences of RNA *in situ*.

Table S2. The primer pairs for dsRNA synthesis.

Table S3. The primer pairs for RT-qPCR detection.

**Figure S1:**
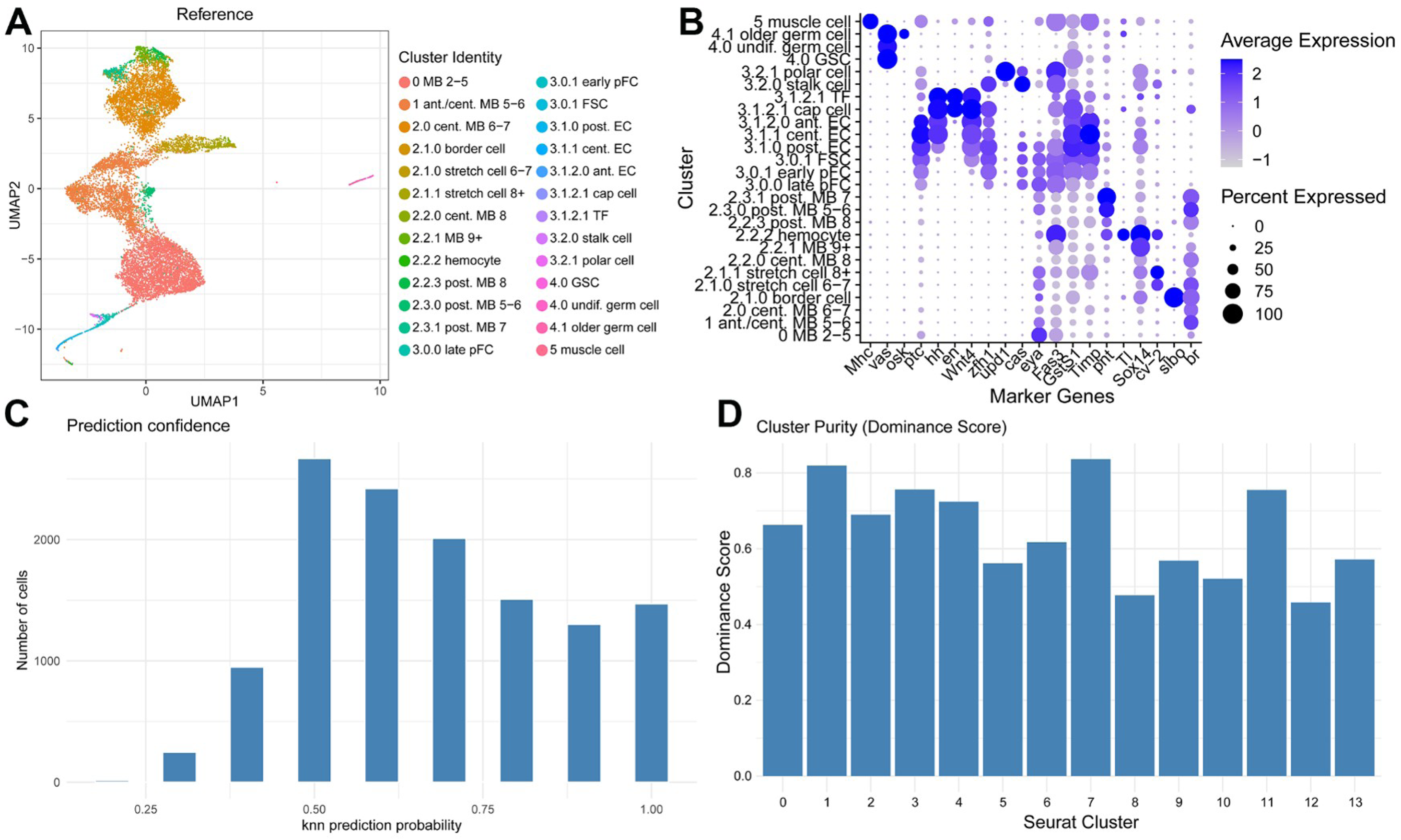
Identification of subclusters in *Drosophila* ovary and prediction probability of cell types in *S. litura* dataset. **A.** A UMAP plot showed different subclusters of referenced *Drosophila* ovary dataset. **B.** Gene expression patterns of marker genes among different cell types in *Drosophila* ovary. **C.** The prediction confidence of cluster identity in *S. litura* dataset. **D.** The cluster purity of 14 clusters was evaluated by Dominance Score.

**Figure S2:**
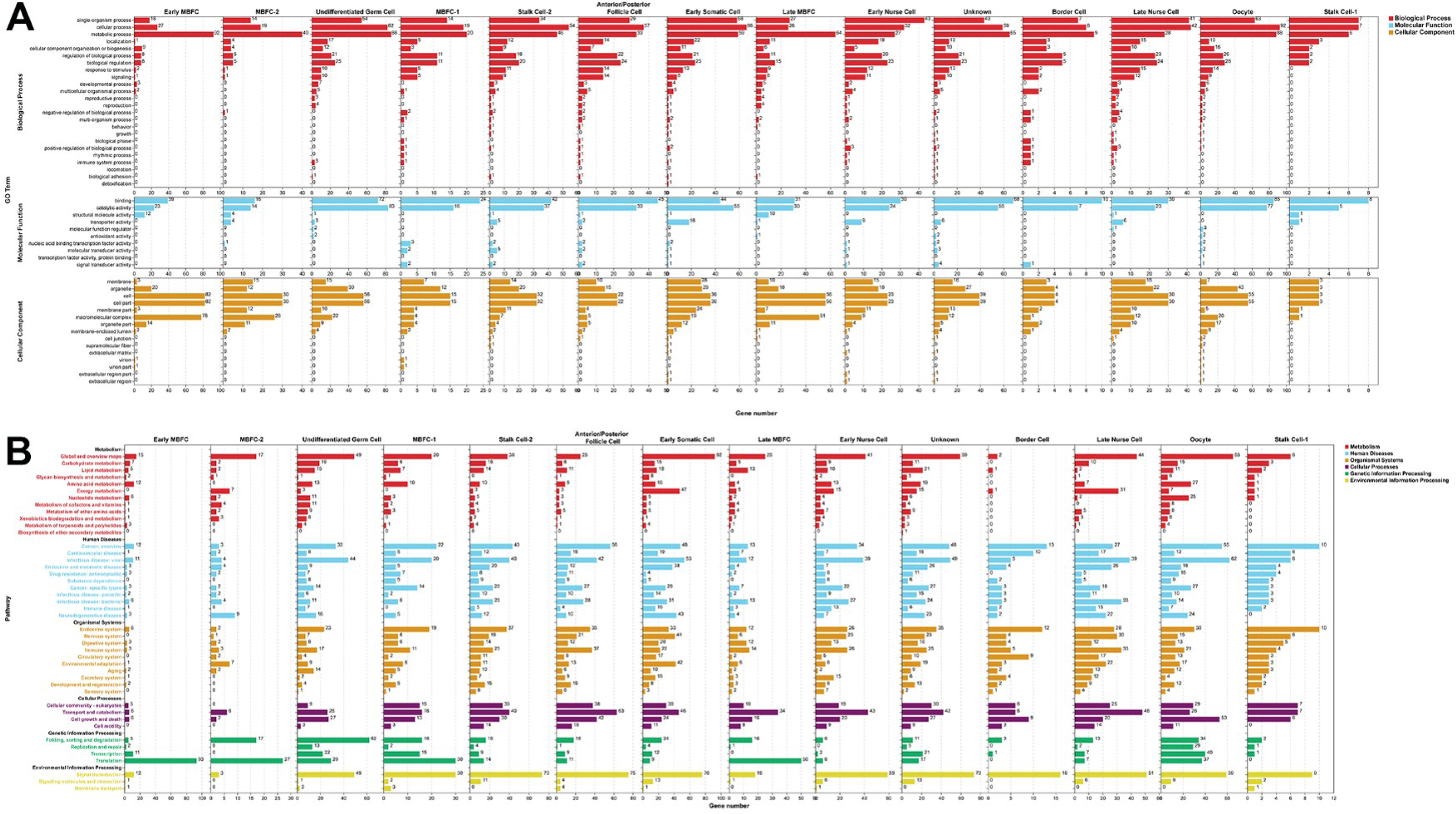
GO/KEGG analysis of each cell type. **A.** GO term analysis was performed for DEGs in each cluster. **B.** KEGG pathway enrichment analysis of each germline cluster and somatic cluster.

